# Physiologically driven, altitude-adaptive model for the interpretation of pediatric oxygen saturation at altitudes above 2000 m a.s.l

**DOI:** 10.1101/334482

**Authors:** Laura Tüshaus, Monica Moreo, Jia Zhang, Stella Maria Hartinger, Daniel Mäusezahl, Walter Karlen

## Abstract

Measuring peripheral oxygen saturation (SpO_2_) with pulse oximeters at the point of care is widely established. However, since SpO_2_ is dependent on ambient atmospheric pressure, the distribution of SpO_2_ values in populations living above 2000 m a.s.l. is largely unknown. Here, we propose and evaluate a computer model to predict SpO_2_ values for pediatric permanent residents living between 0 and 4000 m a.s.l. Based on a sensitivity analysis of oxygen transport parameters, we created an altitude-adaptive SpO_2_ model that takes physiological adaptation of permanent residents into account. From this model, we derived an altitude-adaptive abnormal SpO_2_ threshold using patient parameters from literature. We compared the obtained model and threshold against a previously proposed threshold derived statistically from data and two empirical datasets independently recorded from Peruvian children living at altitudes up to 4100 m a.s.l. Our model followed the trends of empirical data, with the empirical data having a narrower healthy SpO_2_ range below 2000 m a.s.l., but the medians did never differ more than 2.29% across all altitudes. Our threshold estimated abnormal SpO_2_ in only 17 out of 5981 (0.3%) healthy recordings, whereas the statistical threshold returned 95 (1.6%) recordings outside the healthy range. The strength of our parametrised model is that it is rooted in physiology-derived equations and enables customisation. Furthermore, as it provides a reference SpO_2_, it could assist practitioners in interpreting SpO_2_ values for diagnosis, prognosis, and oxygen administration at higher altitudes.

**New & Noteworthy:** Our model describes the altitude-dependent decrease of SpO_2_ in healthy pediatric residents based on physiological equations and can be adapted based on measureable clinical parameters. The proposed altitude-specific abnormal SpO_2_ threshold might be more appropriate than rigid guidelines for administering oxygen that currently are only available for sea level patients. We see this as a starting point to discuss and adapt oxygen administration guidelines.

## INTRODUCTION

Acute lower respiratory infections (ALRI) are a major health burden in low- and middle-income countries. Childhood pneumonia accounts for 14% of all deaths in children worldwide under five years of age (45), of which 95 % occur in low resource settings (41). Common conditions observed in ALRI are dyspnoea and hypoxemia, an abnormally low level of oxygen saturation in the arterial blood (SaO_2_), that can lead to cyanosis and subsequently to death (43). A rapid and non-invasive estimation of hypoxemia can be obtained through pulse oximetry that measures peripheral oxygen saturation (SpO_2_). Pulse oximetry has become a suitable technology for application in low resource settings due to the simplicity of use in combination with mobile phones and non-invasiveness of the device (20, 27). The use of pulse oximeters and supplemental oxygen in clinical applications at the point of care has shown to drastically reduce death rates (8). However, in countries where these devices are needed most, health personnel have only slowly started to gain access.

The interpretation of SpO_2_ values for hypoxemia is challenging, especially for health personnel not familiar with respiratory physiology and measurement principles of pulse oximeters. The World Health Organization (WHO) recommends the administration of oxygen when SpO_2_ drops below or is equal to 90% (44). This fixed threshold oversimplifies hypoxemia treatment (7). It does not provide an indication on when to stop treatment and does not permit adaptation to the local conditions. Namely, in many rural areas, oxygen is a scarce and precious resource and therefore only restrictively administered. Altitude has a direct influence on SpO_2_ as the air pressure decreases, and consequently, the alveolar oxygen partial pressure decreases with increasing altitude (43). Thus, the treatment of ALRI (i.e. administration of oxygen), diagnosis and prognosis, might be affected at higher altitudes and the recommended oxygen administration guidelines at sea level may not be applicable. However, before determining treatment thresholds at higher altitudes, healthy values in this environment need to be established.

In this work, we introduce an altitude-adaptive SpO_2_ model and propose a model-derived altitude-adaptive abnormal SpO_2_ threshold. The physiology-backed altitude-adaptive model describes SpO_2_ values of healthy children living permanently at altitudes up to 4000 m a.s.l. With this model, we aim to provide a better understanding of healthy SpO_2_ values at altitudes above 2000 m a.s.l. for healthy children. The altitude-adaptive abnormal SpO_2_ threshold is obtained by setting the model parameters to abnormal values found in hypoxemic patients. We evaluate these results with a novel dataset obtained from healthy children living in the rural Andes of Peru.

### Related Work

The current literature presents two modelling approaches that describe the relationship between SpO_2_ and altitude.

Subhi et al. developed a statistical model of the SpO_2_ distribution across altitudes that is based on empirical observations from healthy children, and derived an altitude-adaptive threshold for hypoxemia from this model (40). Data were obtained through a literature review of studies performed between 0 and 4018 m a.s.l. A linear random effects meta-regression was performed to predict mean and 2.5^th^ centile SpO_2_ with an exponential equation. This 2.5^th^ centile of healthy children’s SpO_2_ at each altitude was proposed as an altitude-adjusted hypoxemia threshold. It is unclear why this specific, statistically derived threshold was chosen. The obtained statistical model and threshold also did not take other influencing factors, such as measurement protocols, choice of oximeter technology, ethnicity and age range of the studied subjects, into account.

Our group developed a computer model that described the pathway of oxygen throughout the cardio-respiratory body compartments (24, 25). It implemented the oxygen cascade described by West (43). The model used well-established physiological equations to explain how the partial oxygen pressure and oxygen concentrations are interrelated between alveolar gas and peripheral blood (24) (Figure 1). The oxygen cascade describes the oxygen loss from the partial pressure of inspired air to the resulting measurements of SpO_2_ by a pulse oximeter. Therefore, the model was based on physiological parameters and integrated pulse oximeter measurement inaccuracies as reported by the manufacturer. A shortcoming of the model was that it assumed many physiological parameters to be constant and therefore did not consider altitude adaptation. Consequently, it could not correctly describe SpO_2_ measured at higher altitudes, especially in people adapted to these conditions such as permanent residents.

**Figure 1:**
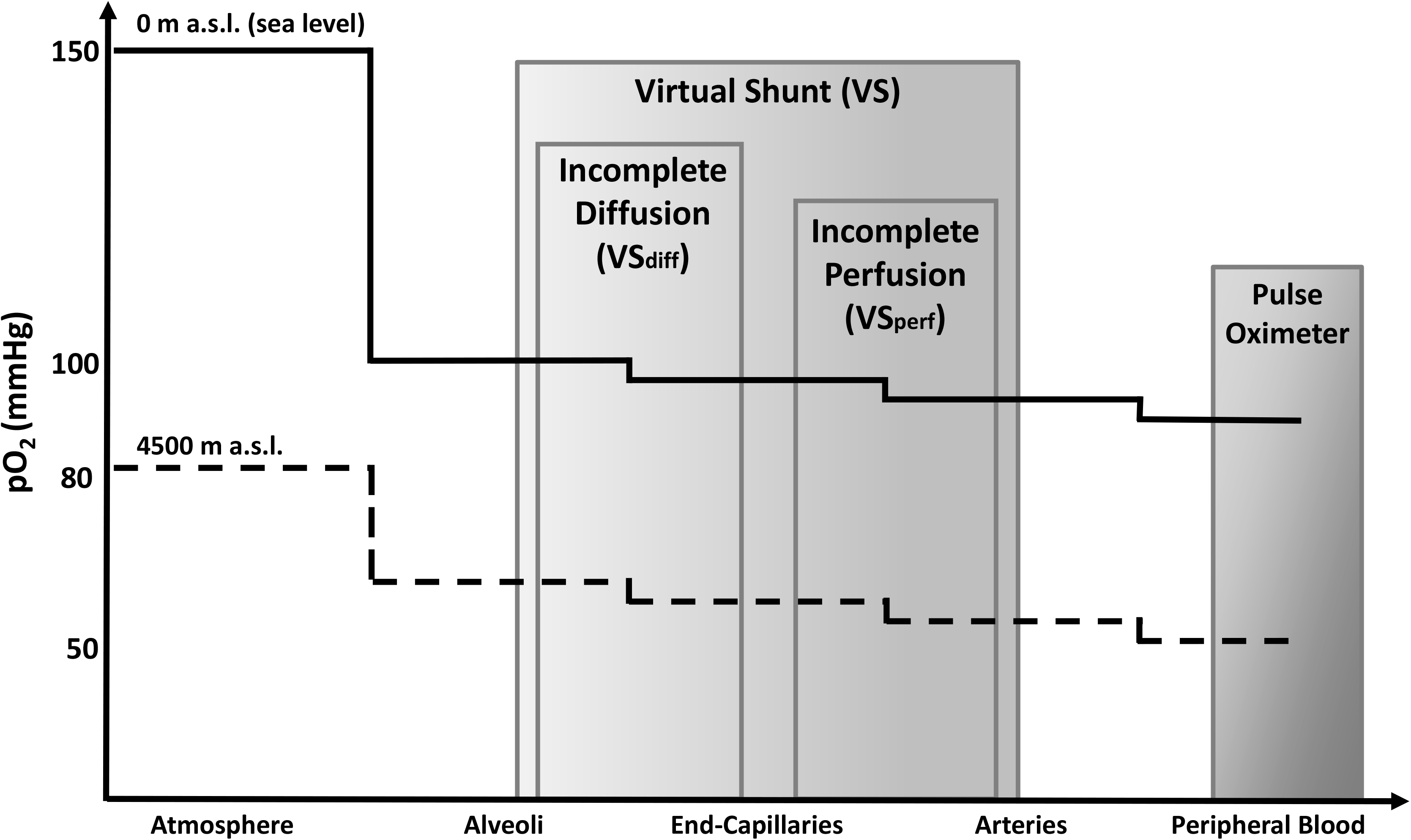
Oxygen cascade describing the loss in oxygen partial pressure (pO_2_) between inspired air and peripheral blood measured with a pulse oximeter. The lines illustrate the standard situation for a healthy subject at sea level (continuous) and at 4500 m a.s.l. (dashed). Virtual shunt describes the combined loss in oxygen content due to incomplete diffusion or perfusion between alveolar and arterial compartments. Adapted from (24, 43).

In a recent prospective study, Rojas-Camayo et al. recorded SpO_2_ from 6289 subjects ranging from infants to elderly people in the Peruvian Andes at 15 altitudes from 154 m to 5100 m a.s.l. (36). They reported the 2.5^th^, 10^th^, 25^th^, 50^th^, 75^th^, 90^th^ and 97.5^th^ centile of the empirical data. This data has not been used to derive a hypoxemia threshold thus far.

## MODELLING

### Altitude-adaptive SpO_2_ model

Starting from the previously established computer model of the oxygen cascade (24), we modified this model to include physiological adaptation to high altitudes. We adjusted parameters that had been found to change with altitude in permanent residents (see Table 1 for an overview of all parameters used). Briefly, the existing model of the oxygen cascade described the pathway of oxygen throughout the cardio-respiratory body compartments (Figure 1) by using physiological equations (see appendix). The model was originally developed to estimate the “virtual shunt” (VS) describing the overall loss of oxygen content between the alveolar gas and arterial blood compartments (2), with SpO_2_ and inspired oxygen (FiO_2_) values as input parameters. An increase in the VS is one of the main causes of hypoxemia (43).

**Table 1:** Physiological parameters obtained from the literature used to implement the altitude-adaptive SpO_2_ model. The healthy ranges (min, max and mean) describe known and expected values in a healthy subject. Parameters that are expected to change under hypoxemic conditions are reported in the last column. We do not differentiate between adults and children. *same value range (min – max) assumed as at sea level. pACO_2_: alveolar partial pressure of carbon dioxide, RQ: respiratory quotient, cHb: haemoglobin concentration, VS_perf_: perfusion defect, VS_diff_: diffusion defect, pAO_2_: alveolar partial pressure of oxygen, FiO_2_: fraction of inspired O_2_, P_atm_: atmospheric pressure, PH_2_O: vapour pressure of water.

| Parameter | Unit | Ref. | Healthy ranges |  |  |  |  |  | Hypoxemia |
| --- | --- | --- | --- | --- | --- | --- | --- | --- | --- |
|  |  |  | Sea level |  |  | 4500 m a.s.l. |  |  |  |
|  |  |  | <u>min</u> | <u>mean</u> | <u>max</u> | <u>min</u> | <u>mean</u> | <u>max</u> |  |
| $pACO_2$ | mmHg | (30, 32, 35) | 35 | 40 | 45 | 23 | 28.3 | 33 | Assumed to be<br>equal to healthy<br>range |
| $RQ$ | unitless | (32, 43) | 0.8 | 0.8 | 1 | 1 | 1 | 1 | |
| $cHb$ | g/100 ml | (43) | 12 | 15 | 17.5 | 17 | 20 | 22.5 | |
| $VS_{perf}$ | % | (31) | 0 | 2 | 5 | 0 | 2 | 5 | $\geq 5^*$ |
| $VS_{diff}$ | % | Derived | 0 | 0 | 0 | 0 | 0 | 0 | $\geq 19^*$ |
| $pAO_2$ | mmHg | Derived | Derived by the alveolar gas equation, with<br>parameters $FiO_2$, $P_{atm}$, $PH_2O$, $pACO_2$ and $RQ$ | | | | | | |

The above mentioned oxygen cascade model, originally developed for adults, can be adapted to a pediatric model as there are no indications that the underlying physics of gas exchange are any different in children (28). We identified relationships between altitude adaptation and parameters of the oxygen cascade, such as atmospheric pressure, haemoglobin concentration (cHb), alveolar partial pressure of carbon dioxide (pACO_2_), and the respiratory quotient (RQ). In addition, we devided VS into two components (Figure 1): 1) incomplete capillary diffusion (diffusion defect between the alveolar and capillary, VS_diff_) and 2) incomplete perfusion with intrapulmonary shunt (perfusion defect, VS_perf_).

We made the following assumptions for the model of a healthy subject: there is no oxygen loss between the alveoli and the end-capillaries (no incomplete capillary diffusion, VS_diff_=0) and SpO_2_ is equal to SaO_2_ (24). These assumptions had the following consequences: the alveolar oxygen partial pressure (pAO_2_) is equal to the partial pressure of oxygen in the end capillaries, the alveolar oxygen saturation is equal to the end capillaries oxygen saturation, and the oxygen content in the alveoli is the same as the in the end capillaries. For the parameters cHb and RQ, we extracted the healthy values at two altitudes (0 m and 4600 m a.s.l.) from the literature (31, 32, 43) and linearly interpolated the parameters between these two altitudes. A linear interpolation was chosen because a sensitivity analysis revealed only small changes upon variation of these parameters (see appendix). For high altitudes (i.e. 4600 m a.s.l.), pACO_2_ was derived from an interpolation of values reported by Rahn and Otis (35), as well as de Meer (32) because the literature presented less coherent values; while for sea level, direct values from Marcdante (30) and West (43) were used. With this information, the oxygen cascade enabled us to estimate the expected SpO_2_ range at a specific altitude. Furthermore, we incorporated the technical tolerances that accounted for the accuracy of pulse oximeters (i.e. ± 2%) determined according to device standards (21) into the model, as shown in Karlen et al. (24). The pulse oximeter accuracy is an important component that is frequently neglected by health practitioners, but influences the pulse oximeter readings and therefore diagnostic results. We include this uncertainty in our model as we strive to better describe the physiology of lung function at different altitudes. Therefore, in the following, when we mention the “healthy ranges”, we refer to the physiological ranges obtained by modelling SpO_2_ based on minimum and maximum literature values of the physiological parameters, combined with the pulse oximeter inaccuracies.

### Altitude-adaptive abnormal SpO_2_ threshold

Analogously, we derive an altitude-dependent threshold for abnormal SpO_2_ by setting model parameters to hypoxemia levels. Hypoxemia is defined as a reduced arterial partial pressure of oxygen (paO_2_), which results in a decrease of SpO_2_ and increase of VS (43). At sea level, as reported in literature, we consider a patient to have hypoxemia if the paO_2_ level is below 80 mmHg (3, 26) and therefore SpO_2_ decreases below 95%. Additionally, we assumed that VS_perf_ increases to above 5%. patients (31). From these assumptions, we recursively derived a disease related increase of VS_diff_ of 19% at sea level. For higher altitudes, we were unable to retrieve any data from the literature that would describe changes (increase or decrease) in VS (VS_diff_ or VS_perf_) or a numerical value for paO_2_ or pAO_2_ under hypoxemia. Therefore, we assumed that the VS components remain constant across altitudes, and the values for cHb, pACO_2_, and RQ are similar in healthy and hypoxemic conditions.

## MATERIALS AND METHODS

To assess the performance and plausibility of our novel altitude-adaptive SpO_2_ model and threshold, we retrospectively evaluated them against a prospectively collected dataset, a previously published dataset, and another, statistical model with threshold.

### Study design and data collection

Our data collection was embedded within a randomised controlled trial by the Swiss-Peruvian Health Research Platform set in the Cajamarca region in the northern highlands of Peru, located in the provinces of San Marcos and Cajabamba. Our study harnessed the operational and logistical setup of this trial, which assessed the efficacy of an Integrated Home-environmental Intervention Package (IHIP-2) to improve child respiratory, enteric, and early development outcomes (19).

The trial was approved by the Universidad Peruana Cayetano Heredia ethical review board and the Cajamarca Regional Health Authority. The trial was registered on the ISRCTN registry (ISRCTN26548981). A total of 317 children aged between 6 and 36 months were enrolled, and informed written consent was obtained from the children’s guardians. A total of 9 field workers (FWs) were trained to visit the children on seven fixed geographical routes. Children were preassigned to these routes and visited in parallel by FWs to perform a mobile health assessment once a week over the course of 60 weeks (6 weeks pilot, followed by a 54-week trial from February 2016 to May 2017, excluding 4 weeks of public holidays). FWs had experience from earlier research projects in collecting basic vital signs and symptoms (17, 18), received five additional days of educational training for the collection of morbidity data, and underwent one month of practical training before the study started (pilot). FWs were equipped with a TAB 2 A7-10 tablet (Lenovo Group Ltd, Beijing, CN). The tablet had a custom mHealth app installed that was developed using the *lambdanative* framework (34). It recorded a photoplethysmogram (PPG) using an USB connected CE marked iSpO_2_ Rx pulse oximeter (Masimo International, Neuchatel, CH) with a multisite Y-probe, and derived SpO_2_ and heart rate (HR). FWs placed the probe on the child’s thumb, index finger, or sole of the foot for the measurement of PPG, HR, and SpO_2_. Simultaneously, respiratory rate was recorded using the RRate app module (22). In addition, the app acquired location and altitude using the embedded global positioning system (GPS) sensor. Furthermore, the app metadata regarding the visit and the recordings such as child ID, timestamps, and child agitation during the vital signs measurements were acquired. All electronically collected data was uploaded from the app into a digital research database (16). Health seeking behaviour and other relevant endpoints were reported in a paper-based, validated questionnaire (18), quality checked, and digitised at the end of the study.

### Post processing

The IHIP-2 vital signs data obtained from the pulse oximeter were post processed to guarantee high data quality. The PPG, SpO_2_, HR, and perfusion index (PI, indication of signal strength) time series from the main trial period were imported into Matlab (R2017b, MathWorks Inc., Natick, USA) where a signal quality index (SQI) for the PPG was calculated (23). We segmented the recordings into segments with SQI > 45. Segments with lower quality (SQI ≤ 45) and with no computed SpO_2_ were excluded. Furthermore, entire recordings were excluded if a single segment duration was shorter than 12 s or the combined length of remaining segments was shorter than 15 s, the range (5^th^ - 95^th^ centile) of SpO_2_ exceeded 5%, and the HR range surpassed 20 bpm in combination with a low perfusion (mean PI ≤ 0.8). We also excluded SpO_2_ values below 60% as they are rare and typically associated with severe clinical cyanosis (46), which was clearly absent in the IHIP-2 cohort. These values also fall in a range where the performance of the pulse oximeters used were not specified by the manufacturer (70% to 100%). Additionally, as each child was always scheduled to be measured weekly at the same altitude (i.e. at home), we verified the consistency of the altitude provided by the GPS. We excluded recordings that contained no altitude information, and altitude outliers that were more than three scaled median absolute deviations away from the median altitude of each child. Altitude outliers could have occurred because at home measurements were not always possible, and because GPS altitude estimates were dependent on weather, the number of available satellites, and other factors. Finally, we excluded measurements which were recorded following a healthcare center visit or the presence of cardio-respiratory or diarrheal disease symptoms in the week preceding the recording. For each remaining high quality recording, we reported the median SpO_2_ over the combined segments of a measurement and the median altitude per child, which was then used for the analysis.

### Evaluation

#### Model

To compare our model with the available datasets, we visualised the altitude dependence of SpO_2_. We applied a locally weighted scatterplot smoothing (lowess) function (5) to all SpO_2_-altitude data pairs collected during the IHIP-2 trial. We limited the comparison to the range of available data (2000–4000 m a.s.l.) to avoid extrapolation errors. Instead of the LMS method used by Rojas-Camayo et al. (36), we reported the centiles of their data with a lowess smoother to ensure equivalent processing of both datasets. Furthermore, we computed the deviations from interpolated medians of both empirical data sets to the model median for each altitude expressed as percent of the respective model value and reported the mean, minimum and maximum deviations. Additionally, we calculated the absolute range of SpO_2_ values at each altitude for both the model and the empirical data sets and reported mean, minimum and maximum range.

#### Threshold

To visualise the differences between the hypoxemia/abnormal SpO_2_ thresholds and oxygen administration guidelines that have been proposed, we graphically compared the altitude-adaptive abnormal SpO_2_ threshold, the statistical hypoxemia threshold, and the WHO guideline for oxygen administration (90%) with the 2.5^th^ centile (lowess smoothed) data of children 1 to 5 years old reported by Rojas-Camayo et al. (36). We further computed the number of measurements in the healthy IHIP-2 data that would have been wrongly classified as abnormal (false positives) when using either the altitude-adaptive abnormal SpO_2_ threshold or the statistical hypoxemia threshold. The false positives are children that are healthy, but likely would receive additional medical attention due to the low SpO_2_ reading.

## RESULTS

We obtained an altitude-adaptive computer model to describe the expected SpO_2_ range in healthy children at higher altitudes, and based on this model proposed a threshold for an abnormal range that could indicate hypoxemia. The parameters used in the mathematical description of the model to define healthy and abnormal ranges are available in Table 1. Out of the 12634 SpO_2_ measurements obtained from 310 children over the course of a year, we retained 5981 measurements from 297 children that were considered complete (contained both GPS and PPG data), featured good quality PPG data, reasonable SpO_2_ (> 60 %) and were recorded when no respiratory disease symptoms or other health issues were reported (410 recordings). At the study start, the mean age of the children was 20.5 months (SD 6.2 months, range: 6-36 months). Each child contributed to a mean of 20.1 (SD 9) repeated measurements. Twenty-one children lived above 3000 m a.s.l. and 8 above 3500 m a.s.l. (Table 2). Therefore, a total of 392 (6.6%) measurements above 3000 m a.s.l. were available.

**Table 2:** IHIP-2 pulse oximeter data: Distribution of children, number of measurements and SpO_2_ per altitude. Age range of the children at study start: 6-36 months (mean: 20.5 months, SD: 6.2 months), 21 children above 3000 m a.s.l. (392 measurements, 6.6 % of total number of measurements), 8 above 3500 m a.s.l. (171 measurements, 2.9 % of total number of measurements), mean number of measurement per child: 20.1 (SD 9).

| Altitude<br>(m a.s.l.) | No.<br>children | Age at study<br>start<br>(months) |  | No. of<br>measurements | No. of measurements per<br>child |  |  | SpO<sub>2</sub> (%) |  |  |
| --- | --- | --- | --- | --- | --- | --- | --- | --- | --- | --- |
|  |  | Mean | SD |  | Min | Mean | Max | Min | Median | Max |
| 2000 | 78 | 21.0 | 6.0 | 1668 | 1 | 21.4 | 39 | 88.9 | 97.2 | 100.0 |
| 2100 | 37 | 20.4 | 6.1 | 803 | 1 | 21.7 | 43 | 87.4 | 97.2 | 100.0 |
| 2200 | 6 | 15.7 | 5.0 | 109 | 7 | 18.2 | 24 | 88.6 | 96.6 | 99.7 |
| 2300 | 15 | 19.5 | 6.7 | 389 | 5 | 25.9 | 34 | 89.2 | 96.7 | 100.0 |
| 2400 | 3 | 24.0 | 2.0 | 77 | 19 | 25.7 | 31 | 91.9 | 96.2 | 100.0 |
| 2500 | 13 | 20.5 | 5.7 | 264 | 4 | 20.3 | 33 | 87.1 | 96.0 | 100.0 |
| 2600 | 22 | 21.6 | 5.6 | 472 | 2 | 21.5 | 39 | 81.1 | 96.0 | 100.0 |
| 2700 | 39 | 21.5 | 6.8 | 753 | 4 | 19.3 | 32 | 85.2 | 95.6 | 100.0 |
| 2800 | 20 | 21.8 | 5.7 | 316 | 1 | 15.8 | 34 | 83.9 | 95.2 | 99.5 |
| 2900 | 18 | 17.7 | 6.0 | 276 | 1 | 15.3 | 27 | 85.6 | 94.7 | 98.6 |
| 3000 | 25 | 20.2 | 7.2 | 462 | 5 | 18.5 | 37 | 81.5 | 94.5 | 100.0 |
| 3100 | 8 | 20.0 | 6.5 | 111 | 1 | 13.9 | 24 | 80.7 | 94.4 | 100.0 |
| 3200 | 3 | 19.3 | 6.0 | 76 | 12 | 25.3 | 38 | 85.7 | 93.4 | 98.2 |
| 3300 | 1 | 20.0 | 0.0 | 19 | 19 | 19.0 | 19 | 89.9 | 92.4 | 95.7 |
| 3400 | 1 | 17.0 | 0.0 | 15 | 15 | 15.0 | 15 | 86.0 | 92.9 | 95.4 |
| 3500 | 0 | - | - | 0 | - | - | - | - | - | - |
| 3600 | 5 | 19.4 | 4.7 | 107 | 9 | 21.4 | 30 | 84.3 | 92.7 | 98.7 |
| 3700 | 1 | 28.0 | 0.0 | 24 | 24 | 24.0 | 24 | 86.4 | 92.5 | 95.6 |
| 3800 | 1 | 23.0 | 0.0 | 22 | 22 | 22.0 | 22 | 80.8 | 87.8 | 93.9 |
| 3900 | 1 | 14.0 | 0.0 | 18 | 18 | 18.0 | 18 | 81.6 | 88.4 | 90.6 |
| **total** | 297 | 20.5 | 6.2 | 5981 | - | 20.1 | - | - | - | - |

### Model

Our altitude-adaptive model provided a SpO_2_ of 97.4% at sea level with a healthy range between 93.5 % and 100% SpO_2_ (Figure 2 and Table 3, high resolution data including model available at (11)). The SpO_2_ of the model decreased with increasing altitude to 89.6% at 4000 m a.s.l. with a healthy SpO_2_ range from 82.3% to 94.1%. The 2.5^th^ and 97.5^th^ centiles reported by Rojas-Camayo et al. largely followed the same trend as those acquired in the IHIP-2 trial, but had a smaller absolute range (Figure 2). Up to 3800 m a.s.l., the 2.5^th^ centiles of both empirical data sets were entirely within the lower boundary of the altitude-adaptive SpO_2_ model’s proposed healthy range, whereas at higher altitudes above 3800 m a.s.l., the 2.5^th^ centile of the IHIP-2 data slightlyfell below this lower boundary. The upper boundary of the altitude-apdaptive SpO_2_ model’s healthy range followed the IHIP-2 data 97.5^th^ centile closely, while it was slightly exceeded by the 97.5^th^ centile data from Rojas-Camayo et al. between 1500 and 3100 m a.s.l by up 0.5 %. In particular, the model showed absolute ranges very similar to both empiricial lowess filtered data sets (model: mean absolute SpO_2_ range: 8.66%, min: 6.42%, max: 11.78%; IHIP-2: mean absolute SpO_2_ range: 8.75%, min: 6.75%, max: 11.22%; Rojas-Camayo: mean absolute SpO_2_ range: 5.53%, min: 3.43%, max: 8.92%). Furthermore, the model differed very little from the interpolated median of the empirical data sets (IHIP-2, deviation of model in percent: mean deviation: 1.5%, min: 0.01%, max: 2.29%; Rojas-Camayo: mean deviation: 1.51%, min: 0.02%, max: 1.95%).

**Figure 2:**
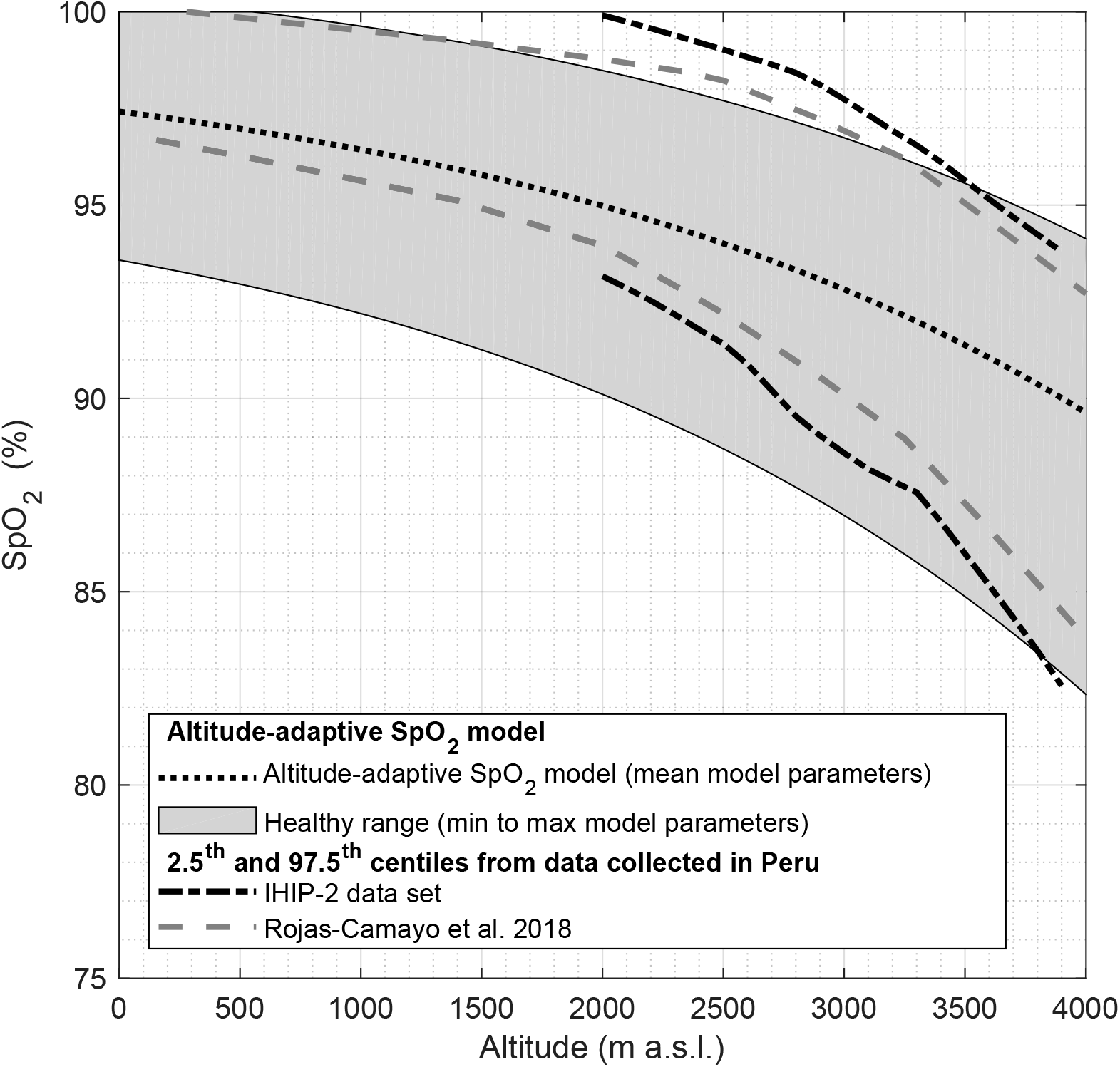
The proposed altitude-adaptive SpO_2_ model provides a healthy SpO_2_ range (light grey area). The black dotted line indicates the median SpO_2_ estimated by the model. The parameters for the min, max and mean model parameters are given in Table 1. The 2.5^th^-97.5^th^centiles of the SpO_2_ data from Rojas-Camayo et al. (light grey dashed lines) (36) and the Integrated Home-environmental Intervention Package (IHIP-2) data set (black dashed-doted lines) that were both recorded in the Peruvian Andes mostly fall into our proposed healthy range. The reported number of measurements per children for the IHIP-2 data can be found in Table 2.

**Table 3:** Values of the healthy ranges of the altitude-adaptive SpO_2_ model including its median and the abnormal SpO_2_ threshold, per altitude. More granular altitude steps available in (11).

| Altitude<br>(m a.s.l.) | Upper healthy range<br>(% SpO<sub>2</sub>) | Lower healthy range<br>(% SpO<sub>2</sub>) | Model Mean<br>(% SpO<sub>2</sub>) | Threshold<br>(% SpO<sub>2</sub>) |
| --- | --- | --- | --- | --- |
| 0 | 100.0 | 93.6 | 97.4 | 92.9 |
| 500 | 100.0 | 93.0 | 97.0 | 92.2 |
| 1000 | 99.6 | 92.2 | 96.4 | 91.2 |
| 1500 | 99.1 | 91.3 | 95.8 | 90.1 |
| 2000 | 98.5 | 90.1 | 95.0 | 88.8 |
| 2500 | 97.7 | 88.7 | 94.0 | 87.1 |
| 3000 | 96.7 | 87.0 | 92.8 | 85.2 |
| 3500 | 95.6 | 84.9 | 91.4 | 82.9 |
| 4000 | 94.1 | 82.3 | 89.6 | 80.1 |

### Threshold

The altitude-adaptive abnormal SpO_2_ threshold followed a similar pattern as the 2.5^th^ centile of Rojas-Camayo’s empirical data with 88.8% vs 94% at 2000 m a.s.l. and 80.1% vs 83.8% at 4000 m a.s.l. (Figure 3, see also Table 3). The 2.5^th^ centile threshold explored by Subhi et al. had an SpO_2_ of 92.8% at 2000 m a.s.l. and then rapidly diverged towards much lower SpO_2_ values for higher altitudes (75.4% at 4000 m a.s.l.). When comparing the two thresholds and their performance for our empirical dataset, the altitude-adaptive threshold estimated abnormal SpO_2_ in only 17 out of 5981 (0.3%) healthy recordings, whereas the 2.5^th^ centile threshold explored by Subhi et al. returned 95 (1.6%) false positives.

**Figure 3:**
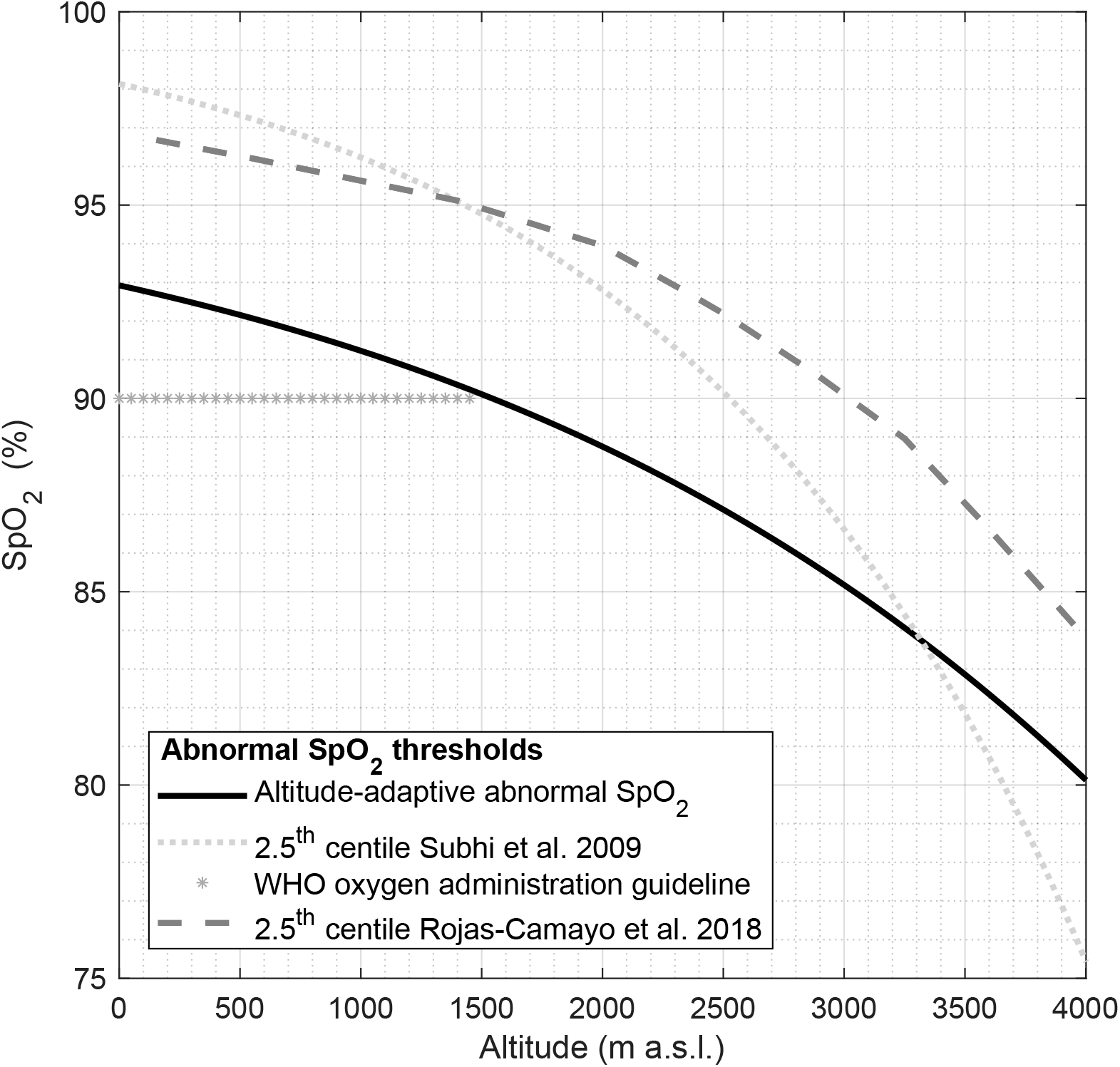
Comparison of proposed abnormal SpO_2_ thresholds that would lead to oxygen administration in patients and existing guidelines. The altitude-adaptive abnormal SpO_2_ threshold (continuous black line) is based on the physiological model derived in this work where a virtual shunt was applied. The threshold from Subhi et al. (40) is the 2.5^th^ centile derived from observations in healthy children collected in a literature review (dotted light grey line), and the 2.5^th^ centile from Rojas-Camayo et al. (36) is derived from a prospectively collected healthy pediatric sample in the Peruvian Andes (dashed grey line). The WHO 90% oxygen administration guideline is a result of a working group consensus (starred grey line) (44) that is in use at lower altitudes.

## DISCUSSION

We proposed an altitude-adaptive model that estimates a healthy SpO_2_ range for children living permanently at altitude and have shown that this proposed healthy SpO_2_ range matches empirical data recorded from a pediatric population living in the Andes. From this model, we derived an altitude-adaptive threshold for abnormal SpO_2_ values. The diagnosis of pneumonia and other respiratory diseases is challenging at altitude, as the most common diagnostic criteria, such as the respiratory rate and oxygen saturation, are dependent on altitude. Our work contributes towards making the management of childhood pneumonia, one of the major causes of child mortality in low resource settings, more objective by attempting to better describe healthy changes of respiratory physiology found in adapted residents. Equipping health workers with mobile pulse oximeters has become an affordable solution, is being evaluated at a large scale (29), and has potential for improving pneumonia treatment at a reasonable cost (12). However, the measurement and interpretation of SpO_2_ can be complicated. Computerised assistance and interpretation of the measurements could ensure reliability of these measurements and provide a meaningful decision support tool to health workers at the central and peripheral level. The proposed adaptive, physiology-based model could provide a basis for the necessary computations because it provides a reference for healthy values at higher altitudes.

Our model is unique as the adjustment of the parameters can be tuned individually, based either on measurements or on known parameter ranges, and it is based on physiology. It was developed considering, where available, literature-based physiological parameter values of Peruvian Andes residents that are adapted to this environment. These parameters could be adjusted without altering the underlying model for other populations with known differences in genetic or physiological adaptation mechanisms (e.g. Himalayan residents) (1).

In contrast to our parameterized model, Subhi and colleagues fitted empirical data collected from across the world into a statistical model describing the SpO_2_ distribution using centiles (40). The statistical model was built using aggregated data collected from mixed populations using pulse oximeters with partially unknown specifications. The statistical model therefore cannot be adjusted to factors such as population-specific variations or varying technical specifications (e.g. differing accuracy of pulse oximeter brands or types). In relation to the two empirical data sets mentioned in this publication, and in comparison to our proposed abnormal SpO_2_ threshold, the statistical threshold provided a very sensitive cut-off at lower altitudes (up to 3300 m a.s.l.). However, it underestimates potentially abnormal SpO_2_ values at higher altitudes. Most likely, this underestimation of the abnormal SpO_2_ values at higher altitudes is due to less data samples being available for the statistical modeling. Our physiological model was not affected by data sparsity, which is a distinctive feature and clear advantage at higher altitudes. Both model thresholds, and the studied data sets, supported the current WHO constant threshold of 90% SpO_2_ for oxygen administration at altitudes below 1500 m a.s.l.

The altitude-adaptive model described the SpO_2_ ranges observed from the empirical data sets with highly similar mean absolute ranges., However, the two empirical datasets presented in this work originate solely from the Peruvian Andes and a single type of pulse oximeter. To further validate the model, it will be crucial to apply data from other regions and ethnicities, and establish if a customised model is required when used in different parts of the world. Such data collection should be accompanied by a gold standard, such as blood gas measurements with information on cHb, SaO_2_, paO_2_ and paCO_2_, in order to pinpoint the exact sources of potentially observed differences.

At higher altitudes above 3800 m a.s.l., we notice higher deviations in the model compared to what is seen in the empirical data due to a slower decline of SpO_2_ in the model. We suspect that this is directly linked to the assumptions we made during the modelling of healthy ranges. We assumed that cHb and RQ change linearly with altitude. However, the adaptation process is likely more pronounced at higher altitudes (6) and might contribute to non-linear parameter changes.

Our assumptions to define the abnormal physiological parameters could limit the validity of the abnormal threshold. We only based our assumptions on literature values that referred to sea level patients. Due to the underlying changes in physiology caused by adaptation, disease manifestation and progression, symptoms could be different at high altitudes compared to at sea level. Furthermore, it is unclear if comorbidities that have not been captured in the present modelling, such as malnutrition, iron deficiency, or diarrheal diseases that are known to negatively influence outcomes of patients with pneumonia (4, 37, 39), would also influence the model parameters. Additional empirical data of sick children are needed to establish models that describe the dependence of these parameters to altitude. For example, anaemic children display altered ranges for blood gas parameters and their actual health status is not entirely captured through our cardio-respiratory model based on SpO_2_ measurements. SpO_2_ and derived hypoxemia estimations reflect only the proportion of O_2_ that is bound to Hb and not the total O_2_ carrying capacity and concentration. Consequently, pulse oximeter assessments are blind to the effective O_2_ available in the tissues. Also, cardiac output, an alternative path to modulate O_2_ delivery (14), is not easily obtainable with pulse oximetry alone. Thus, clinicians need to take the overall clinical situation of the child into consideration and evaluate treatment options accordingly when interpreting hypoxemia thresholds (10).

To assess the performance of the model, we limited the comparison to altitudes from 2000 to 4000 m a.s.l. where corresponding empirical data was available. The data contained weekly measurements for each child repeated over a full year (mean: 20.1, SD: 9), therefore representing the expected measurement and physiological variability within a healthy subject. Among the children recruited from the Cajamarca region during the IHIP-2 trial, only 21 lived above 3000 m a.s.l. which increases the variability in the data. Nevertheless, we observed very similar SpO_2_ ranges from Rojas-Camayo et al. (36). Despite the high numbers of repeated measurements and rigid measurement protocols, both datasets showed a high variability in the measured SpO_2_. For example, in the IHIP-2 dataset, at 2000 m a.s.l. a healthy range corresponded to 11% (Table 2). Our model represented this large range of possible healthy values accurately. Nevertheless, the inter- and intra-individual variability could originate from a number of sources not incorporated in the model. Circadian variation in pediatric SpO_2_ has been reported (42) and we did not account for such daytime differences. Furthermore, there are known sex differences in adults (1), which could also apply to the pediatric population. Although we used the most recent pulse oximeter technology and performed continuous measurements for at least a minute with a rigorous approach to PPG post-processing for high quality, not all the sources for measurement errors in pulse oximetry, such as poor perfusion, inacurate probe positioning, or ambient light interference (13), could be fully excluded in this dataset.

Additionally, it is important to note that neonates were not considered in the modeling process. Neonatal blood is known to benefit from the high affinity of fetal haemoglobin and would have changed the oxygen dissociation curve considerably (33). Since hyperoxia in neonates leads to oxidative stress with potentially severe health complications (15), the definition of an abnormal threshold and consequently the guideline for oxygen administration would require a more detailed, separate discussion for this population.

We established an altitude-adaptive abnormal SpO_2_ threshold based on physiologically plausible values. Our results show that using such a threshold is most relevant at altitudes above 2000 m a.s.l. The 90% SpO_2_ threshold recommended by the WHO for oxygen administration in patients living at sea level clearly does not apply to these altitudes. Compared to the previously published statistical altitude-dependent threshold by Subhi et al. (40), our threshold leads to fewer detections of false positives (healthy children falsely categorized as hypoxemic). Conversely, while Subhi et al. also promoted the use of an altitude-dependent threshold at higher altitudes (2500 m a.s.l.), their threshold is very conservative at altitudes below 2950 m a.s.l. but more lenient at higher altitudes, where it decreases very steeply which might exclude a number of patients in need of supplemental oxygen.

### Outlook

Thus far, experts have not agreed on a definition for abnormal SpO_2_ thresholds at altitudes higher than sea level. To date, no reliable SpO_2_ data from children suffering from hypoxemia and ALRI at altitude are available. The advancement of research for developing better tools to diagnose pneumonia and ALRI at altitude would greatly benefit from access to publicly available, comprehensive data sets obtained from sick children.

With pulse oximeters increasingly being used as monitors for ALRI diagnosis and treatment, additional research is urgently needed to provide a reliable description of the SpO_2_ distribution at altitude, and to develop guidelines of oxygen administration for hypoxemic children living in these settings.

Furthermore, knowledge of abnormal SpO_2_ values at high altitudes could help in the development of new decision support tools for health workers operating in low resource settings with the goal to improve clinical management of hypoxemia in children with ALRI in the future.

## CONCLUSION

Improvement of SpO_2_-altitude models present a first step towards an integration of pulse oximetry in low resource settings and could further the development of valid altitude-dependent thresholds for treatment of childhood pneumonia and other ALRI. We developed an altitude-adaptive physiology-backed SpO_2_ model using an existing physiological model using the concept of VS adjusted for published ranges of values for pACO_2_, cHb, and RQ. Based on this model, healthy ranges and an altitude-dependent abnormal SpO_2_ threshold are suggested that are based on physiological variations of vital parameters. With the increased availability of sensors and digitalised systems in low resource settings, parametrised models could provide additional valuable support to primary health workers to understand the patient’s condition at the point of care, and choosing treatment options based on objectively obtained physiological measurements.

## ACKNOWLEDGMENTS

We are grateful to all staff and students from the Swiss – Peruvian Health Research Platform and the San Marcos research station, especially Angelica Fernandez and Maria Luisa Huyalinos, Hector Verastegui and Nestor Nuño for their assistance and support throughout the study. The San Marcos Red Salud-IV health personnel supported the SpO_2_ measurement in the peripheral health posts. Matthias Hüser programmed the assessment app and maintained the software throughout the study. We would like to thank all the families that participated in the randomised trial. We thank Dr. Jose Rojas-Camayo for sharing the centiles of his valuable dataset, Ms Janine Burren for her valuable input on statistics and data representation, and Dr. Urs Frey and Joanne Lim for helpful comments on this manuscript. We appreciate the various contributions of the colleagues from the Swiss Pediatric Surveillance Unit (SPSU) network. Furthermore, Masimo International kindly facilitated the access to their pulse oximeter sensors in Peru.

## Appendix

**Table A1:**
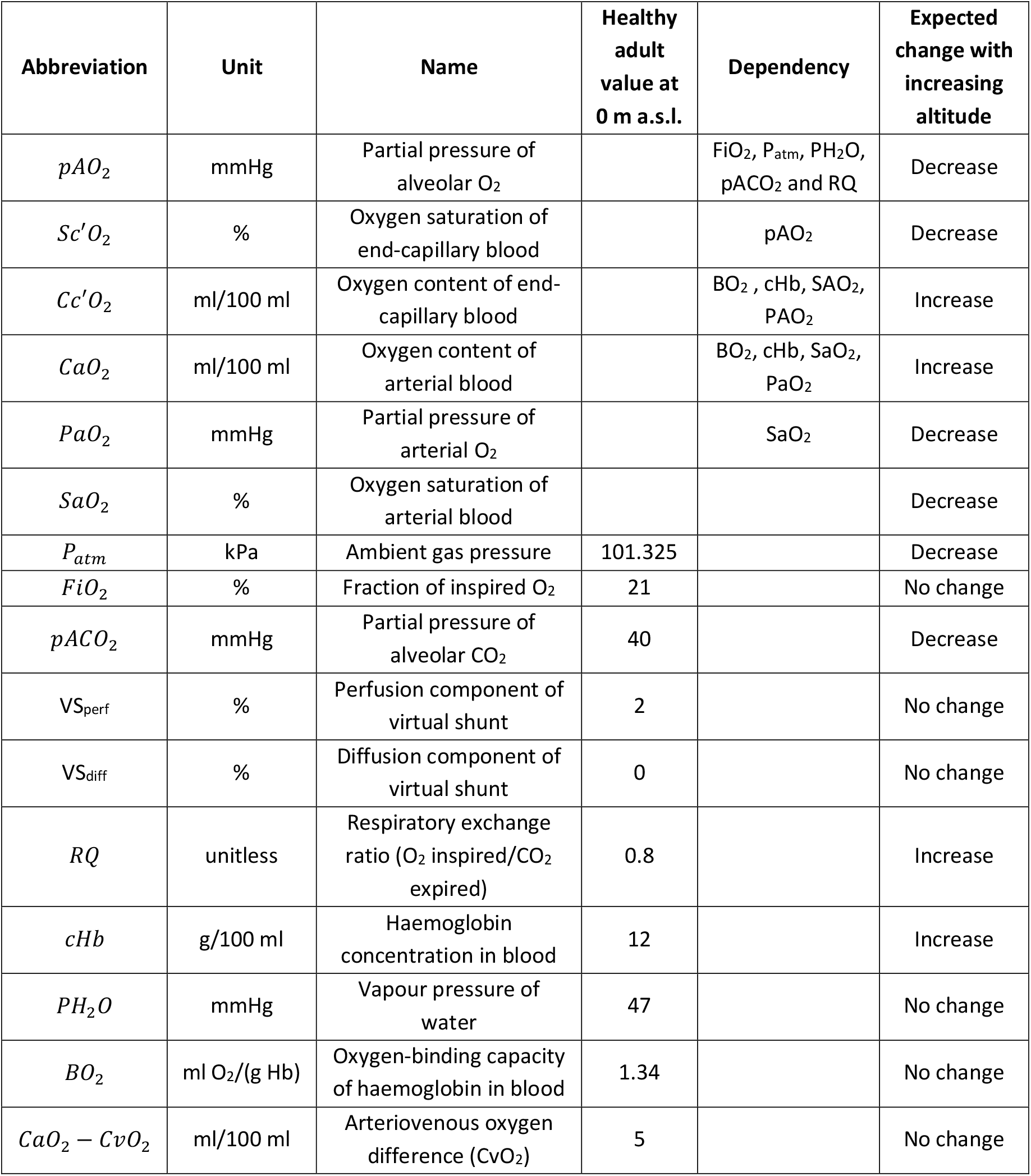
Parameters required for the calculation of the oxygen cascade with specification of dependency on other parameters and the expected change with increasing altitude.

### Equations

For the entire computer model of the oxygen cascade, please consult (24, 25). See Table A1 for the variable names.

*Alveolar gas equation:*

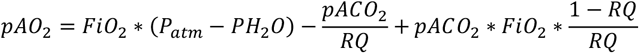

*Severinghaus equation (38):*

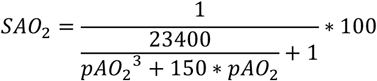

*O_2_ Content equation:*

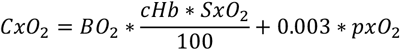

*Severinghaus-Ellis equation (9):*

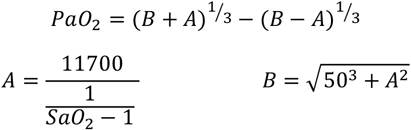

*Virtual Shunt from perfusion defect (VS_perf_):*

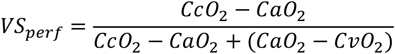

*Virtual Shunt from diffusion defect (VS_diff_):*

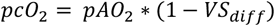

### Sensitivity analysis

To display the influence of parameters on the output of the oxygen cascade, a sensitivity analysis was performed (Figure A1). The parameters variation was chosen to reproduce the minimum and maximum value used in the altitude-adaptive SpO_2_ model (Table 1). A change in pACO_2_ had the highest effect, followed by VS_diff_, VS_perf_ and RQ. A change in cHb is negligible for the calculation of SpO_2_, however, please note that it has a significant influence on availability of O_2_ in the tissues.

**Figure A1:**
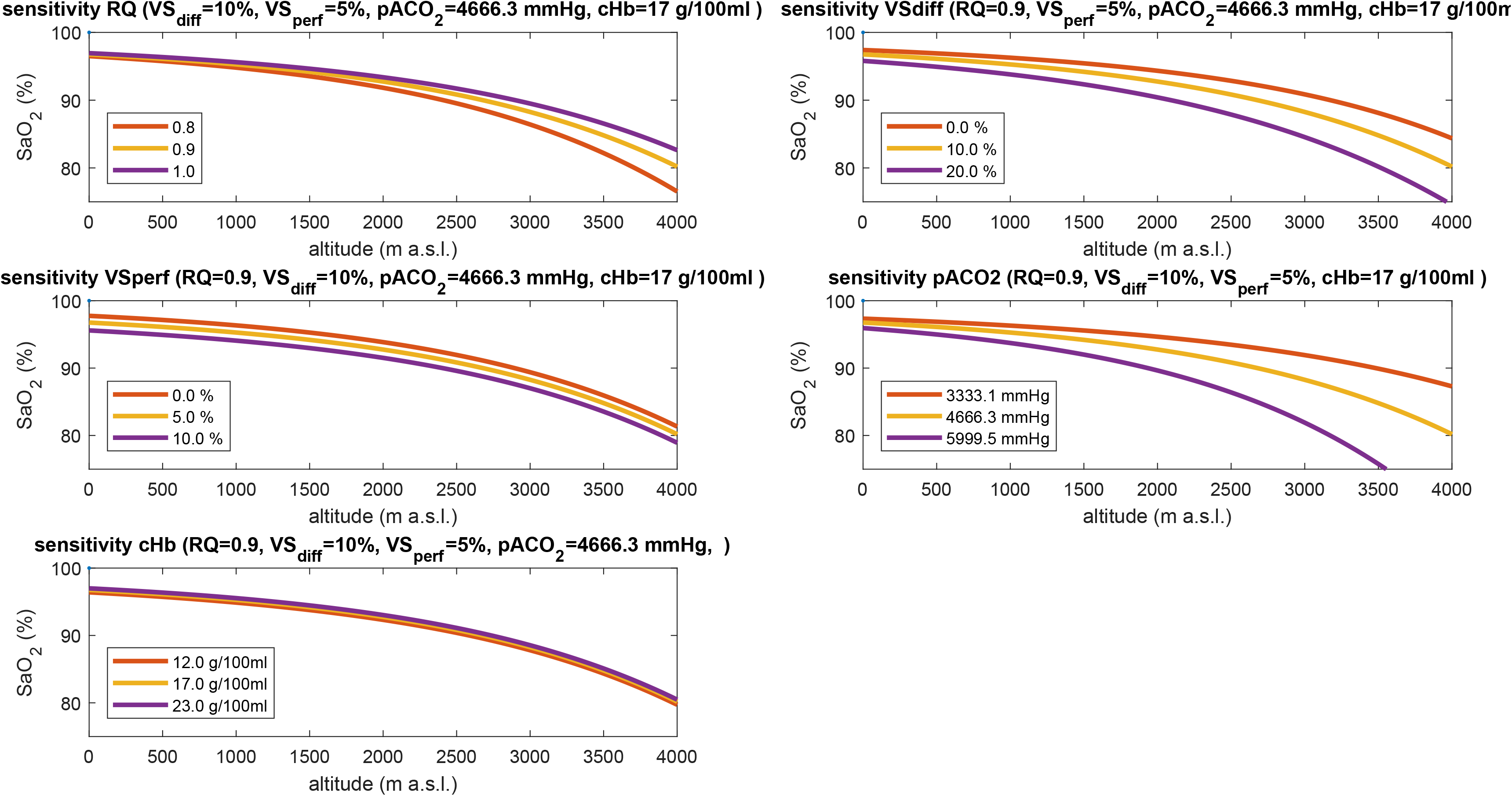
Sensitivity analysis of the five main model parameters respiratory quotient (RQ) (top left), virtual shunt from diffusion defect (VS_diff_)(top right), virtual shunt from perfusion defect (VS_perf_) (middle left), PACO_2_ (middle right), and cHb (bottom left).

## REFERENCES

1. Beall CM. Two routes to functional adaptation: Tibetan and Andean high-altitude natives. Proc Natl Acad Sci 104: 8655–60, 2007.

2. Benatar SR, Hewlett AM, Nunn JF. The use of ISO-shunt lines for control of oxygen therapy. Brit J Anaesth 45: 711–8, 1973.

3. Bradley JS, Byington CL, Shah SS, Alverson B, Carter ER, Harrison C, Kaplan SL, MacE SE, McCracken GH, Moore MR, St Peter SD, Stockwell JA, Swanson JT. The management of community-acquired pneumonia in infants and children older than 3 months of age: Clinical practice guidelines by the pediatric infectious diseases society and the infectious diseases society of America. Clin Infect Dis 53: 25–76, 2011.

4. Chisti MJ, Graham SM, Duke T, Ahmed T, Faruque ASG, Ashraf H, Bardhan PK, Shahid ASMSB, Shahunja KM, Salam MA. Post-discharge mortality in children with severe malnutrition and pneumonia in Bangladesh. PLoS One 9: 1–7, 2014.

5. Cleveland WS, Devlin SJ. Locally Weighted Regression: An Approach to Regression Analysis by Local Fitting. J Am Stat Assoc 1459: 37–41, 1988.

6. Cohen JH, Haas JD. Hemoglobin correction factors for estimating the prevalence of iron deficiency anemia in pregnant women residing at high altitudes in Bolivia. Rev Panam Salud Pública/Pan Am J Public Heal 6: 392–9, 1999.

7. Duke T, Subhi R, Peel D, Frey B. Pulse oximetry: technology to reduce child mortality in developing countries. Ann Trop Paediatr 29: 165–75, 2009.

8. Duke T, Wandi F, Jonathan M, Matai S, Kaupa M, Saavu M, Subhi R, Peel D. Improved oxygen systems for childhood pneumonia: a multihospital effectiveness study in Papua New Guinea. Lancet 372: 1328–33, 2008.

9. Ellis RK. Determination of PO2 from saturation. J Appl Physiol 67: 902, 1989.

10. Enoch AJ, English M, Shepperd S. Does pulse oximeter use impact health outcomes? A systematic review. Arch Dis Child 101: 694–700, 2016.

11. ETH Zurich Research Collection. Altitude-adaptive model for pediatric oxygen saturation. (2019). doi: 10.3929/ethz-b-000344084.

12. Floyd J, Wu L, Hay Burgess D, Izadnegahdar R, Mukanga D, Ghani AC. Evaluating the impact of pulse oximetry on childhood pneumonia mortality in resource-poor settings. Nature 528: S53–9, 2015.

13. Fouzas S, Priftis KN, Anthracopoulos MB. Pulse Oximetry in Pediatric Practice. Pediatrics 128: 740–52, 2011.

14. Gutierrez JA, Theodorou AA. Oxygen Delivery and Oxygen Consumption in Pediatric Critical Care. In: Pediatric Critical Care Study Guide, edited by Lucking SE, Maffei FA, Tamburro RF, Thomas NJ. Springer, p. 19–38.

15. Habre W, Peták F. Perioperative use of oxygen: Variabilities across age. Br J Anaesth 113: ii26–36, 2014.

16. Harris PA, Taylor R, Thielke R, Payne J, Gonzalez N, Conde JG. Research electronic data capture (REDCap) - a metadata-driven methodology and workflow process for providing translational research informatics support. J Biomed Inform 42: 377–81, 2009.

17. Hartinger SM, Lanata CF, Hattendorf J, Gil AI, Verastegui H, Ochoa T, Mäusezahl D. A community randomised controlled trial evaluating a home-based environmental intervention package of improved stoves, solar water disinfection and kitchen sinks in rural Peru: Rationale, trial design and baseline findings. Contemp Clin Trials 32: 864–73, 2011.

18. Hartinger SM, Lanata CF, Hattendorf J, Verastegui H, Gil AI, Wolf J, Mäusezahl D. Improving household air, drinking water and hygiene in rural Peru: A community-randomized-controlled trial of an integrated environmental home-based intervention package to improve child health. Int J Epidemiol 45: 2089–99, 2016.

19. Hartinger SM, Nuno N, Hattendorf J, Verastegui H, Ortiz M, Mäusezahl D, Nuño N, Verastegui H, Ortiz M, Mäusezahl D. A factorial randomised controlled trial to combine early child development and environmental interventions to reduce the negative effects of poverty on child health and development: rationale, trial design and baseline findings. BioRxiv (2018). doi: 10.1101/465856.

20. Hudson J, Nguku SM, Sleiman J, Karlen W, Dumont GA, Petersen CL, Warriner CB, Ansermino JM. Usability testing of a prototype phone Oximeter with healthcare providers in high- and low-medical resource environments. Anaesthesia 67: 957–67, 2012.

21. International Standard Organisation. ISO 80601-2-61 Medical electrical equipment — Part 2-61: Particular requirements for basic safety and essential performance of pulse oximeter equipment. Geneva: 2011.

22. Karlen W, Gan H, Chiu M, Dunsmuir D, Zhou G, Dumont GA, Ansermino JM. Improving the accuracy and efficiency of respiratory rate measurements in children using mobile devices. PLoS One 9: e99266, 2014.

23. Karlen W, Kobayashi K, Ansermino JM, Dumont GA. Photoplethysmogram signal quality estimation using repeated Gaussian filters and cross-correlation. Physiol Meas 33: 1617–29, 2012.

24. Karlen W, Petersen CL, Dumont GA, Ansermino JM. Variability in estimating shunt from single pulse oximetry measurements. Physiol Meas 36: 967–81, 2015.

25. Karlen W, Petersen CL, Dumont GA, Mark Ansermino J. Corrigendum: Variability in estimating shunt from single pulse oximetry measurements. Physiol Meas 38: 1746–7, 2017.

26. Kasper D, Fauci AS, Hauser SL, Longo DL, Jameson JL, Loscalzo J, editors. Harrison’s Principles of Internal Medicine. 19th ed. New York, USA: McGraw-Hill Education, 2015.

27. Luks A, Swenson E. Pulse oximetry at high altitude. High Alt Med Biol 12: 109–19, 2011.

28. Madden K, Khemani RG, Newth CJL, Argent AC, Hatherill M, Newth CJ, Klein M, Barker SJ, Tremper KK, Barker SJ, Tremper KK, Hyatt J, Corkey CW, Barker GA, Edmonds JF, Al. E, Frey B, Butt W, Gagnon S, Jodoin A, Martin R, Hammer J, Newth CJ, Hammer J, Eber E, Hedstrand U, Hoover CF, Kim YJ, Lenke LG, Bridwell KH, Al. E, Lenke LG, Bridwell KH, Blanke K, Baldus C, Maniscalco M, Zedda A, Faraone S, Al. E, Mithoefer JC, Bossman OG, Thibeault DW, Mead GD, Newth CJL, Hammer J, Nichols MM, Nichols DG, Numa AH, Newth CJ, Perez A, Mulot R, Vardon G, Al. E, Permutt S, Rayner J, Trespalacios F, Machan J, Al. E, Sivan Y, Deakers TW, Newth CJ, Sivan Y, Eldadah MK, Cheah TE, Newth CJ, Stalcup SA, Mellins RB, Steele DW, Santucci KA, Wright RO, Al. E, Tobias JD, Flanagan JF, Wheeler TJ, Al. E, West JB, Zhang JG, Wang W, Qiu GX, Al. E. Paediatric applied respiratory physiology – the essentials. Paediatr Child Health (Oxford) 19: 249–56, 2009.

29. Malaria Consortium. The Pneumonia Diagnostics Project: evaluating devices for accuracy [Online]. 2016. https://www.malariaconsortium.org/resources/publications/739/The-Pneumonia-Diagnostics-Project:-evaluating-devices-for-accuracy-[5Jun.2019].

30. Marcdante KJ, Kliegman RM, editors. NELSON Essentials of Pediatrics. 7th ed. Philadelphia, USA: Elsevier Saunders, 2015.

31. Marino PL, editor. The ICU Book. 4th ed. Philadelphia, USA: Lippincott Williams & Wilkins, 2013.

32. de Meer K, Heymans HSA, Zijlstra WG. Physical adaptation of children to life at high altitude. Eur J Pediatr 154: 263–272, 1995.

33. Nelson NM, Prodhom LS, Cherry RB, Smith CA. A Further Extension of the in Vivo Oxygen-Dissociation Curve for the Blood of the Newborn Infant. J Clin Invest 43: 606–10, 1964.

34. Petersen CL, Gorges M, Dunsmuir D, Ansermino M, Dumont GA. Experience report: Functional Programming of mHealth Applications. In: Proc. of the 18th ACM SIGPLAN int conference on Functional programming. ACM Press, p. 357–62.

35. Rahn H, Otis AB. Man’s respiratory response during and after acclimatization to high altitude. Am J Physiol 157: 445–62, 1949.

36. Rojas-Camayo J, Mejia CR, Callacondo D, Dawson JA, Posso M, Galvan CA, Davila-Arango N, Bravo EA, Loescher VY, Padilla-Deza MM, Rojas-Valero N, Velasquez-Chavez G, Clemente J, Alva-Lozada G, Quispe-Mauricio A, Bardalez S, Subhi R. Reference values for oxygen saturation from sea level to the highest human habitation in the Andes in acclimatised persons. Thorax 87: 1–4, 2017.

37. Schlaudecker EP, Steinhoff MC, Moore SR. Interactions of diarrhea, pneumonia, and malnutrition in childhood. Curr Opin Infect Dis 24: 496–502, 2011.

38. Severinghaus JW. Simple, accurate equations for human blood O2 dissociation computations. J Appl Physiol 46: 599–602, 1979.

39. Singh V, Aneja S. Pneumonia - management in the developing world. Paediatr Respir Rev 12: 52–9, 2011.

40. Subhi R, Smith K, Duke T. When should oxygen be given to children at high altitude? A systematic review to define altitude-specific hypoxaemia. Arch Dis Child 94: 6–10, 2009.

41. United Nations Children’s Fund (UNICEF). Pneumonia and diarrhoea: Tackling the deadliest diseases for the world’s poorest children. New York, USA: United Nations Children’s Fund (UNICEF), 2012.

42. Vargas MH, Heyaime-Lalane J, Pérez-Rodríquez L, Zúñiga-Vázquez G, Furuya MEY. Day-night fluctuation of pulse oximetry: An exploratory study in pediatric inpatients. Rev Investig Clin 60: 303–10, 2008.

43. West JB. Respiratory Physiology: the essential. 9th ed. Baltimore, USA: Lippincott Williams & Wilkins, 2012.

44. World Health Organization. Pocket book of hospital care for children: Second edition Guidelines for the management of common childhood illnesses. 2nd ed. Geneva, CH: World Health Organization, 2013.

45. World Health Organization, The United Nations Children’s Fund (UNICEF). Ending preventable child deaths from pneumonia and diarrhoea by 2025: The integrated Global Action Plan for Pneumonia and Diarrhoea (GAPPD). Geneva, CH: World Health Organization/The United Nations Children’s Fund (UNICEF), 2013.

46. Yadav A, editor. Monitoring in Anesthesia. In: Short Texbook of Anesthesia. 2018, p. 61.

